# Top2a-dependent neuronal regulation of social behavior and persistent rescue of social deficit through PRC2-mediated epigenetic reprogramming

**DOI:** 10.64898/2026.02.24.707855

**Authors:** Baixi He, Yuxuan Mao, Catherine Hong, Yijie Geng

## Abstract

Disruption of social behavior is a core feature of autism spectrum disorder (ASD), yet the molecular and cellular mechanisms governing social behavior development are not well understood. We previously identified topoisomerase IIα (Top2a) as a critical regulator of social behavior and restricted and repetitive behavior through antagonism of polycomb repressive complex 2 (PRC2)-mediated H3K27 trimethylation (H3K27me3), based mainly on pharmacological perturbations during embryogenesis in zebrafish and mouse. However, whether neuronal Top2a is genetically required for social behavior in mammals, and whether PRC2 inhibition can rescue genetically induced social deficits, remain untested. Here, we establish a neuron-specific Top2a conditional knockout mouse model and demonstrate that neuronal Top2a haploinsufficiency selectively impairs social interaction without inducing restricted and repetitive behaviors or cognitive deficits. Pharmacological inhibition of PRC2 using the EZH2 inhibitor tazemetostat, combined with elacridar to facilitated blood-brain barrier penetration, robustly rescues social deficits in Top2a conditional knockout mice. Strikingly, a one-week oral dosing regimen produced a rescue effect that persisted for up to two months after treatment cessation, far exceeding the temporal window typically observed for neuromodulatory drugs targeting neurotransmitter systems. These results showcase the unique capability of epigenetic modulatory therapy to induce durable behavioral improvements and their therapeutic potential for treating social dysfunction in neuropsychiatric disorders. Together, our results provide direct genetic evidence that neuronal Top2a governs social behavior in mice and establish the neuronal Top2a-PRC2 axis as a conserved, targetable epigenetic pathway regulating social behavior.

## INTRODUCTION

Social behavior is a fundamental and evolutionarily conserved feature of animal life, enabling cooperation, communication, and social learning. In humans, early-emerging social behaviors^1^ form the foundation for later cognitive and emotional development^2,3^, while disruptions in social interaction represent a core and defining feature of autism spectrum disorder (ASD). Although hundreds of genetic variants contributing modest risk for ASD have been identified^4,5^, how these diverse genetic and environmental factors converge on shared biological pathways that regulate social behavior development remains poorly understood.

Topoisomerase IIα (Top2a) is a DNA-modifying enzyme best known for its essential roles in DNA replication, transcription, and genome organization. Despite its classification as a ubiquitous “housekeeping” enzyme, accumulating evidence suggests that Top2a may play a selective role in neurodevelopment and ASD-related biology. Large-scale human genetic analyses have identified TOP2A as a hub gene within autism-associated gene networks^6,7^, and transcriptional studies have implicated Top2a in regulating long, neurodevelopmentally important genes enriched for ASD risk^8-10^. However, whether Top2a plays a causal and cell-type-specific role in regulating social behavior in mammals has remained unresolved.

In previous work, we used a chemical-genetic screening approach in zebrafish to identify Top2a inhibition during embryogenesis as a potent disruptor of social behavior^11^. Transient embryonic exposure to structurally diverse Top2a inhibitors induced persistent social deficits in zebrafish and mice, accompanied by selective downregulation of ASD risk genes. In addition to social deficits, prenatal Top2a inhibition also induced restricted and repetitive behaviors in mice. Mechanistically, these effects were linked to aberrant regulation of polycomb repressive complex 2 (PRC2)-mediated H3K27 trimethylation (H3K27me3). Pharmacological inhibition of EZH2, the core catalytic subunit of PRC2, was sufficient to rescue social deficits in zebrafish. These findings identified a conserved Top2a-PRC2 epigenetic pathway regulating social behavior development and suggested a potential mechanistic link between environmental Top2a inhibition and ASD risk^11^.

Despite these advances, several critical questions remained unanswered. First, because prior conclusions were largely based on pharmacological inhibition, especially in mice experiments, it was unclear whether Top2a is genetically required within the mammalian brain for normal social behavior, or whether observed phenotypes reflected indirect effects of chemical exposure. Second, although PRC2 inhibition rescued social deficits in zebrafish, it remained unknown whether epigenetic rescue is effective in a mammalian system, particularly in the context of genetically induced Top2a deficiency. Third, it was unknown whether the two core behavioral symptoms of ASD, namely social deficits and restricted and repetitive behaviors, are regulated by the Top2a-PRC2 pathway through shared or distinct brain regions and cell types. Fourth, it remained unclear whether other ASD-associated behavioral alterations, including cognitive deficits, anxiety-related behavior, and depressive-like behavior, are also regulated by the Top2a-PRC2 pathway. Addressing these questions is essential for establishing the Top2a-PRC2 pathway as a bona fide developmental mechanism rather than a pharmacological artifact and defining the scope and specificity of its ASD-relevant behavioral regulatory effects.

From a therapeutic perspective, most currently available neuropsychiatric drugs act by acutely modulating neurotransmitter systems and typically require continuous administration to maintain behavioral effects^12-16^. Such neuromodulatory interventions rarely produce lasting behavioral improvements after treatment cessation^17,18^. In contrast, epigenetic regulatory mechanisms offer the potential to durably reprogram gene expression states underlying behavior^19-21^. Whether transient pharmacological modulation of epigenetic pathways can induce long-lasting rescue of genetically encoded social deficits in mammals, however, remains largely unexplored. Demonstrating sustained behavioral recovery following short-term epigenetic intervention would represent a major conceptual and translational advance for the treatment of neurodevelopmental and psychiatric disorders.

Here, we address these gaps by generating and characterizing a neuron-specific Top2a conditional knockout mouse model. We show that neuronal Top2a haploinsufficiency in the brain selectively impairs social interaction while sparing restricted and repetitive behavior, anxiety-like behavior, cognition, and depressive-like behavior, revealing a dissociation between social behavior and other ASD-relevant phenotypes. We further demonstrate that pharmacological inhibition of PRC2 using the FDA-approved EZH2 inhibitor tazemetostat, with brain penetration enhanced by co-administration of the dual ABCB1/ABCG2 inhibitor elacridar^22^, robustly and persistently rescues social deficits in Top2a conditional knockout mice. Together, these findings provide direct genetic validation of neuronal Top2a as a selective regulator of social behavior and establish PRC2 inhibition as an effective strategy to persistently reverse social deficits caused by Top2a deficiency in mammals.

## RESULTS

### Neuron-specific Top2a haploinsufficiency in the brain selectively impairs social interaction in mice

To genetically test whether neuronal Top2a is required for social behavior in mammals, we generated neuron-specific Top2a conditional knockout mice using a Nestin-Cre-driven recombination strategy (Fig. 1A). Both male and female heterozygous Top2a^flx/+^ mice were crossed with homozygous Nestin-Cre mice to produce littermates consisting of Top2a^flx/+^;Nestin-Cre conditional knockout (cKO) mice and Top2a^+/+^;Nestin-Cre wild-type controls. Genotypes were confirmed by PCR prior to behavioral testing. A series of behavioral assays were carried out to evaluate ASD-relevant behavioral phenotypes and control phenotypes, including social interaction (Fig. 1B), exploratory behavior (Fig. 1C: object exploration assay), locomotion and anxiety-like behavior (Fig. 1D: open field assay), restricted and repetitive behaviors (Fig. 1E: T-maze assay; Fig. 1F: marble burying assay), cognition (Fig. 1G: novel object recognition assay), and depressive-like behavior (tail suspension assay). All assays were conducted blinded to genotype.

**Figure 1.**
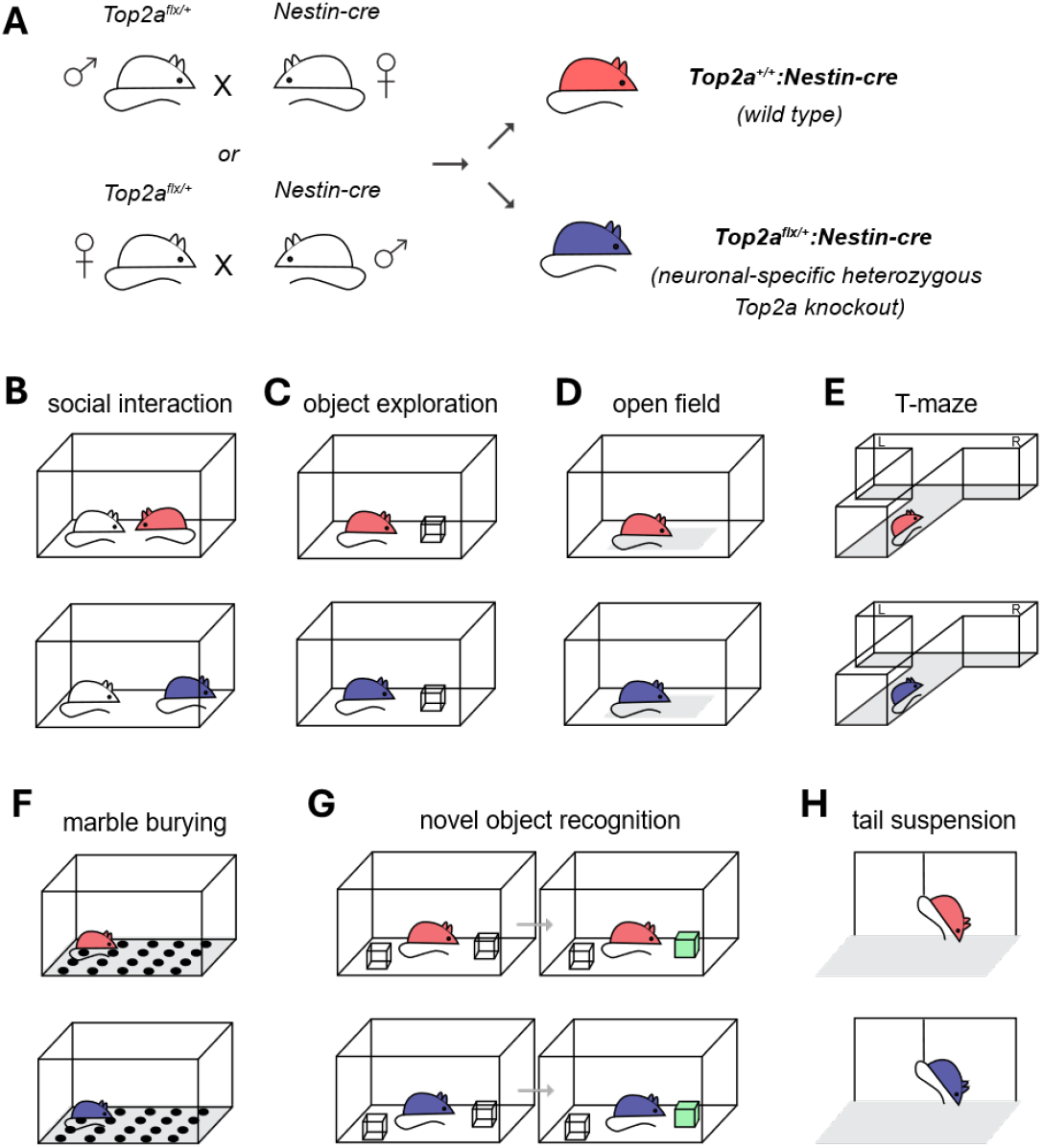
Schematic illustrations of the experimental design for characterizing behavioral impacts of Top2a conditional knockout in neurons. (**A**) Schematic of the breeding strategy. Heterozygous Top2a^flx/+^ mice were crossed with homozygous Nestin-Cre mice to generate Top2a^flx/+^;Nestin-Cre conditional knockout (cKO) mice and Top2a^+/+^;Nestin-Cre wild-type littermate controls. (**B**) Social behavior was assessed using a reciprocal social interaction assay in which a test mouse was placed in a neutral arena with an unfamiliar, age- and sex-matched wild-type conspecific. Social interaction behaviors were quantified through total duration of social interactions, mean duration, and total number of interactions. (**C**) Exploratory behavior was assessed using a novel object exploration assay in which a test subject was placed in a neutral arena with an unfamiliar object. Exploratory behaviors were quantified through total duration of explorations, mean duration, and total number of explorations. (**D**) Locomotor activity and anxiety-like behavior were assessed using the open field assay by measuring exploration of the center versus peripheral zones of the arena. (**E**) Restricted and repetitive behavior was assessed using the T-maze assay, in which mice were allowed to freely choose between left and right arms across repeated trials. (**F**) Restricted and repetitive behavior was assessed using the marble burying assay, in which the number of marbles buried or displaced over a fixed period was quantified. (**G**) Recognition memory was evaluated using the novel object recognition assay, in which mice were exposed to familiar and novel objects and preference for the novel object was quantified. (**H**) Depressive-like behavior was assessed using the tail suspension assay, in which immobility time was quantified during suspension by the tail.

Social behavior was assessed using a reciprocal social interaction assay in which a test mouse was placed in a neutral arena with an unfamiliar, age- and sex-matched wild-type conspecific (Fig. 1B). Compared to wild-type littermates, male neuronal Top2a cKO mice exhibited a significant reduction in total social interaction time, particularly when measuring the total duration (Fig. 2A) and mean duration (Fig. 2B) of social interactions, whereas the total number of social interactions remained comparable between the cKO mice and control mice (Fig. 2C). To determine whether the observed social deficit reflected a sex-specific effect, we analyzed male and female mice separately. Similar to the males, female Top2a cKO mice also showed significantly reduced social interaction compared to their respective wild-type controls when measuring the total duration of their social interactions (Fig. 2D). Mean duration of social interaction was also reduced, although not able to reach statistical significance (Fig. 2E; *p*=0.056). Similar to males, the total number of social interaction was not significantly altered in female Top2a cKO mice compared to the controls (Fig. 2F). Together, these results demonstrate that neuronal Top2a is genetically required for normal social interaction in mice and indicate that neuronal Top2a haploinsufficiency impairs social behavior in a sex-independent manner.

**Figure 2.**
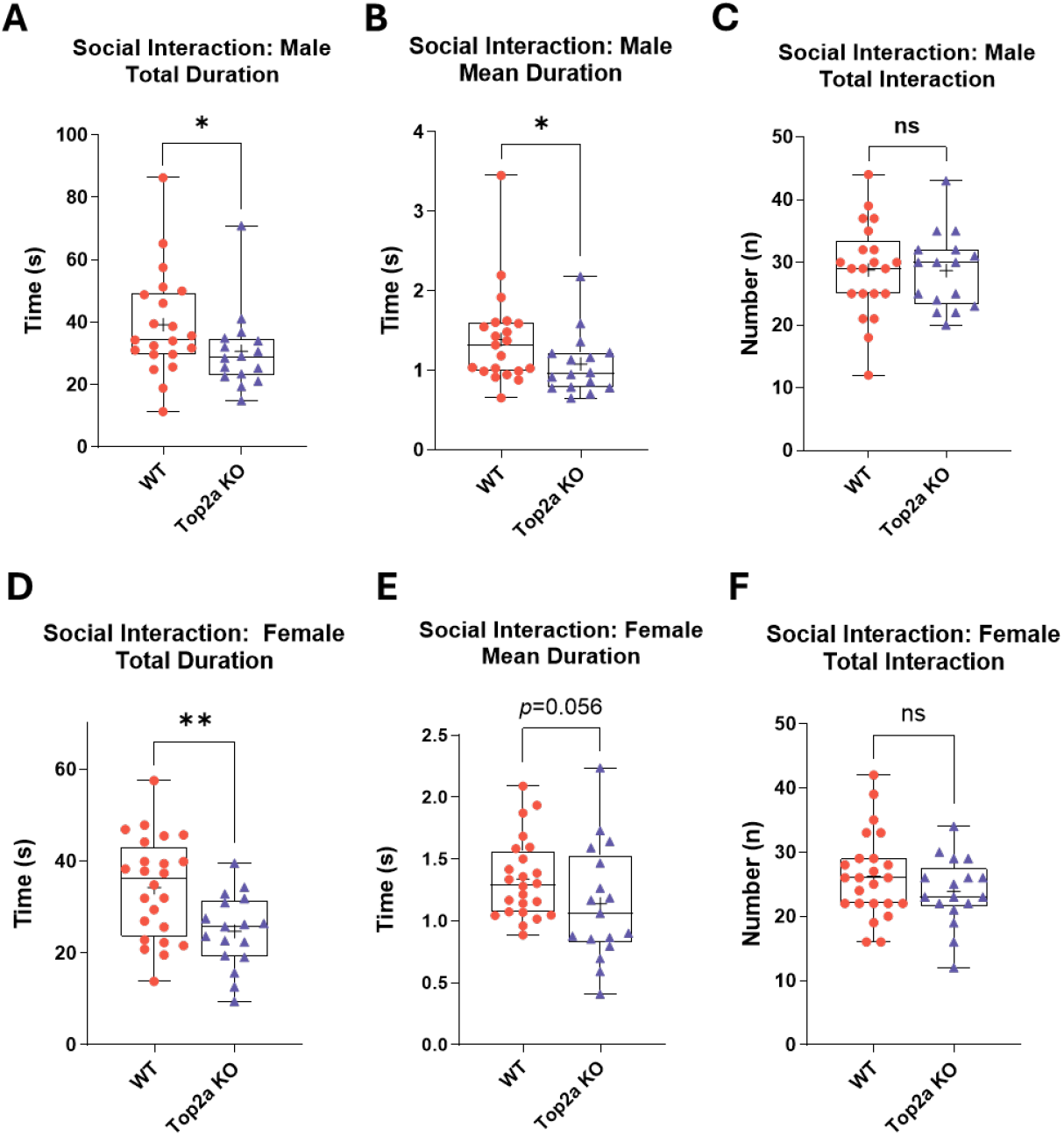
Neuron-specific Top2a haploinsufficiency selectively impairs social interaction in mice. (**A-C**) Social interaction metrics in male mice. Compared to wild-type controls, male Top2a cKO mice showed a significant reduction in total duration of social interaction (A) and mean duration per interaction (B), while the total number of social interactions was not significantly altered (C). (**D-F**) Social interaction metrics in female mice. Female Top2a cKO mice exhibited a significant reduction in total duration of social interaction (D) and a trend toward reduced mean duration per interaction (E), while the total number of social interactions was not significantly different from controls (F). Each dot represents an individual mouse. Box plots indicate median (center line), interquartile range (box), and range (whiskers). All behavioral scoring was performed blinded to genotype. Sample sizes were as follows: male wild-type, *n* = 21; male Top2a cKO, *n* = 16; female wild-type, *n* = 24; female Top2a cKO, *n* = 17. Statistical significance was determined using one-tailed Student’s *t* tests. ns: not significant; *: *p*<0.05, **: *p*<0.01.

To evaluate whether the reduction in social interaction was selective, we examined behavioral parameters that could confound social performance if impaired, including exploratory behavior, locomotion, and anxiety-like behavior. In the novel object exploration assay, male Top2a cKO mice explored the novel object at levels comparable to wild-type controls, as indicated by similar total duration (Fig. 3A), mean duration (Fig. 3B), and number of exploratory bouts (Fig. 3C). In female mice, we observed a modest increase in total exploration time in Top2a cKO animals (Fig. 3D). This effect was not driven by longer individual exploration events (Fig. 3E), but rather by a slight increase in the number of exploratory bouts (Fig. 3F); however, this difference did not reach statistical significance. These results indicate that the social deficits observed in Top2a cKO mice were not attributable to reduced exploratory drive or general behavioral disengagement. General locomotor activity and anxiety-like behavior were further assessed using the open field assay. Both male (Fig. 3G & 3H) and female (Fig. 3I & 3J) Top2a cKO mice exhibited locomotor activity and center exploration comparable to wild-type controls, indicating the absence of locomotor deficits or altered anxiety-like behavior. In fact, female Top2a cKO mice showed a trend toward increased time spent in the center of the open field compared to wild-type controls, although this difference did not reach statistical significance (Fig. 3J). Together, these findings demonstrate that the social deficits induced by neuronal Top2a depletion are selective and cannot be explained by impairments in exploratory behavior, locomotion, or anxiety-related processes.

**Figure 3.**
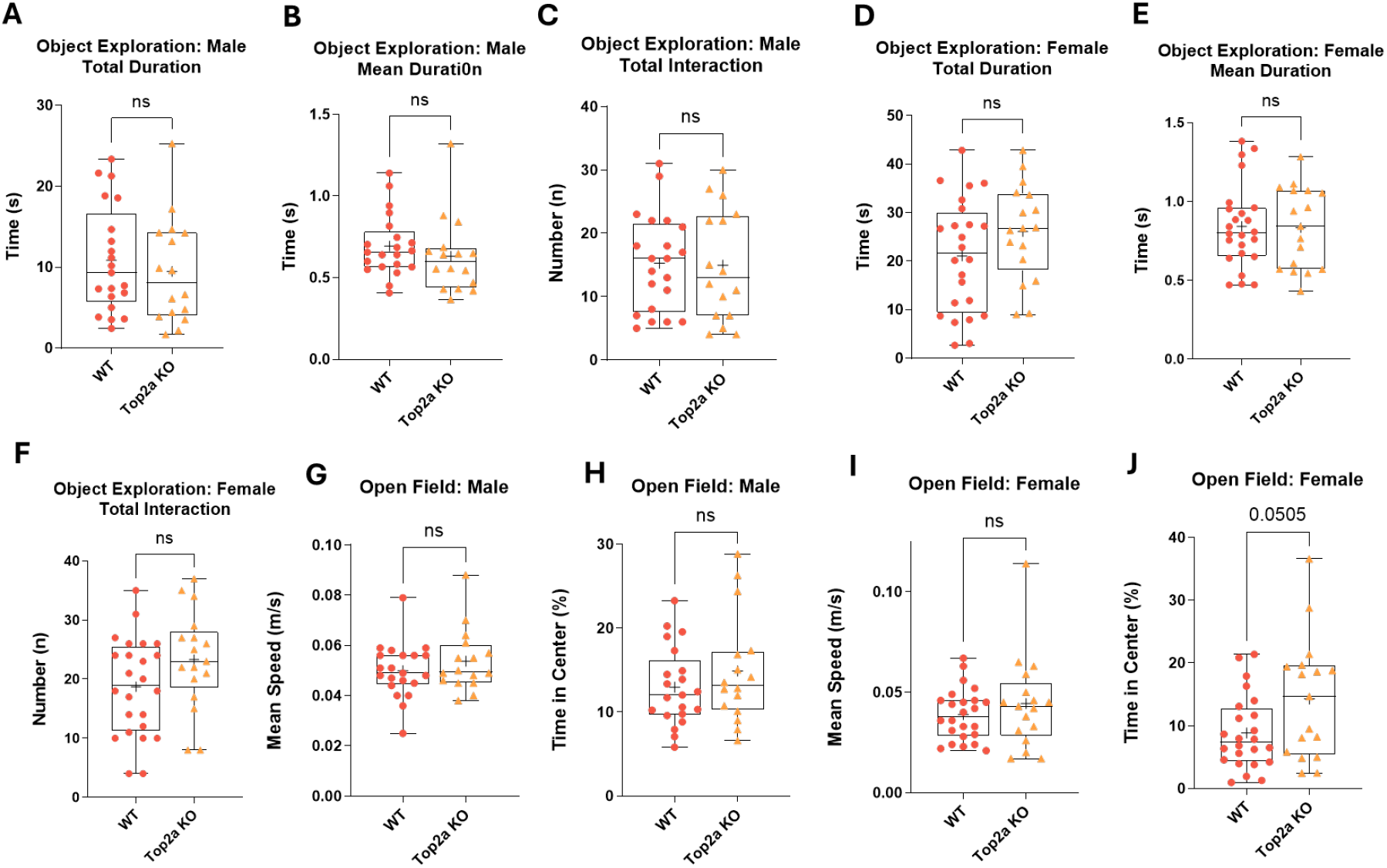
Neuronal Top2a haploinsufficiency does not impair exploratory behavior, locomotion, or anxiety-like behavior. (**A-C**) Novel object exploration behavior in male mice. Top2a conditional knockout (cKO) mice exhibited exploration levels comparable to wild-type controls, as measured by total exploration duration (A), mean exploration duration (B), and number of exploratory bouts (C). (**D-F**) Novel object exploration behavior in female mice. Top2a cKO mice showed a modest, non-significant increase in total exploration duration (D), which was not driven by longer individual exploration events (E) but instead reflected a slight increase in the number of exploratory bouts (F). (**G, H**) Open field assay in male mice showing comparable total distance traveled (G) and time spent in the center zone (H) between Top2a cKO mice and wild-type controls, indicating normal locomotor activity and anxiety-like behavior. (**I, J**) Open field assay in female mice showing no significant differences in total distance traveled (I) or time spent in the center zone (J) between genotypes. Female Top2a cKO mice displayed a trend toward increased center exploration that did not reach statistical significance. Each dot represents an individual mouse. Box plots indicate median (center line), interquartile range (box), and range (whiskers). All behavioral scoring was performed blinded to genotype. Sample sizes were as follows: male wild-type, *n* = 21; male Top2a cKO, *n* = 16; female wild-type, *n* = 24; female Top2a cKO, *n* = 17. Statistical significance was determined using two-tailed Student’s *t* tests. ns: not significant.

### Neuronal Top2a haploinsufficiency does not induce restricted and repetitive, cognitive, or depressive-like behavioral deficits

To determine whether neuronal Top2a haploinsufficiency broadly disrupts ASD-relevant behaviors beyond social interaction, we next assessed restricted and repetitive behaviors, cognitive function, and depressive-like behavior in Top2a conditional knockout mice using a battery of established behavioral assays (Fig. 1E-1H). Restricted and repetitive behaviors was evaluated using the marble burying assay and the T-maze spontaneous alternation task. Both male (Fig. 4A & 4B) and female (Fig. 4C & 4D) Top2a cKO mice displayed marble burying behavior comparable to wild-type littermates, with no significant differences in the number of marbles buried or displaced during the assay. Similarly, performance in the T-maze did not differ between genotypes in either sex, indicating intact exploratory choice behavior and absence of perseverative arm selection (Fig. 4E-4H).

**Figure 4.**
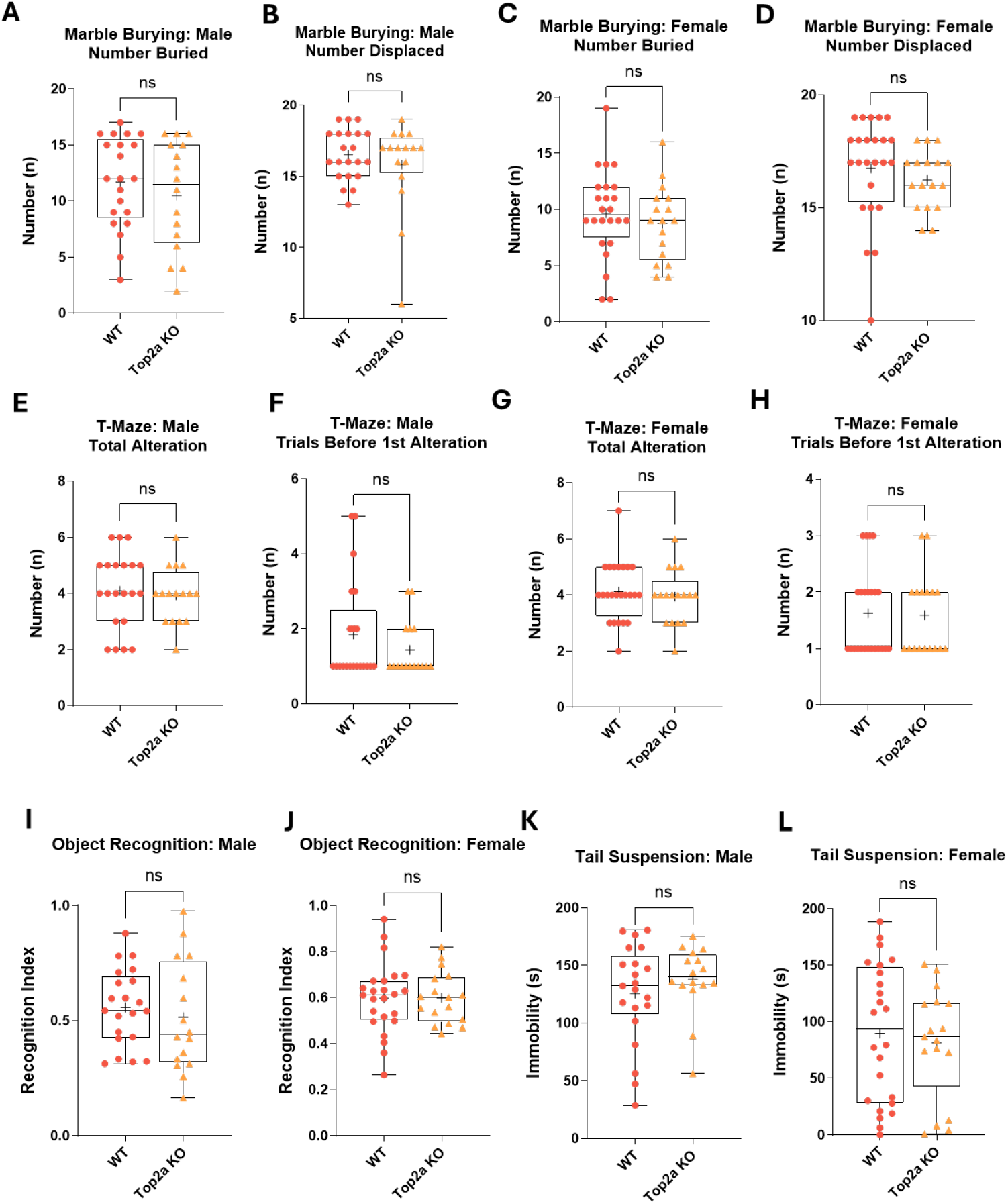
Neuronal Top2a haploinsufficiency does not induce restricted and repetitive, cognitive, or depressive-like behavioral deficits. (**A, B**) Marble burying assay in male mice showing no significant differences between Top2a conditional knockout (cKO) mice and wild-type controls in the number of marbles buried (A) or displaced (B). (**C, D**) Marble burying assay in female mice showing comparable marble burying behavior between Top2a cKO mice and wild-type controls. (**E-H**) T-maze spontaneous alternation task assessing restricted and repetitive behavior. Both male (E, F) and female (G, H) Top2a cKO mice exhibited alternation performance indistinguishable from wild-type littermates, as measured by number of total alterations (E, G) and number of trials before the first alteration event (F, H), indicating intact exploratory choice behavior and absence of perseverative arm selection. (**I, J**) Novel object recognition assay assessing cognitive function. Both male (I) and female (J) Top2a cKO mice showed normal preference for the novel object relative to the familiar object, as quantified by recognition index: time spent investigating the novel object / total time spent investigating novel and familiar objects. (**K, L**) Tail suspension assay assessing depressive-like behavior. Immobility duration was comparable between Top2a cKO mice and wild-type controls in both males (K) and females (L), indicating no induction of depressive-like behavior by neuronal Top2a haploinsufficiency. Each dot represents an individual mouse. Box plots indicate median (center line), interquartile range (box), and range (whiskers). All behavioral scoring was performed blinded to genotype. Sample sizes were as follows: male wild-type, *n* = 21; male Top2a cKO, *n* = 16; female wild-type, *n* = 24; female Top2a cKO, *n* = 17. Statistical significance was determined using two-tailed Student’s *t* tests. ns: not significant.

We next assessed cognitive function using the novel object recognition assay. Both male (Fig. 4I) and female (Fig. 4J) Top2a cKO mice exhibited normal preference for the novel object relative to the familiar object comparable to wild-type controls, as demonstrated by similar recognition index, indicating preserved recognition memory and learning. Depressive-like behavior, a commonly reported comorbidity in ASD, was evaluated using the tail suspension assay. Both male (Fig. 4K) and female (Fig. 4L) Top2a cKO mice exhibited immobility durations comparable to those of wild-type controls, with no significant genotype-dependent differences observed, indicating that neuronal Top2a haploinsufficiency does not induce depressive-like behavioral alterations. Together, these results demonstrate that neuronal Top2a haploinsufficiency selectively impairs social interaction without inducing restricted and repetitive, cognitive, or depressive-like deficits. This behavioral dissociation indicates that neuronal Top2a plays a specific role in regulating social behavior rather than broadly affecting multiple ASD-relevant behavioral domains.

### Pharmacological inhibition of PRC2 sustainably rescues social deficits in neuronal Top2a cKO mice

We next tested whether pharmacological inhibition of PRC2 could rescue the social deficits induced by neuronal Top2a haploinsufficiency in mice. Based on our prior findings in zebrafish and the known role of EZH2-mediated H3K27 trimethylation downstream of Top2a, we treated adult Top2a cKO mice with the EZH2 inhibitor tazemetostat in combination with the dual ABCB1 and ABCG2 inhibitor elacridar to enhance brain penetration^22^ (Fig. 5A). Following a 7-day oral dosing regimen, social behavior was assessed longitudinally after treatment cessation, with the final day of oral gavage designated as 0 day post-treatment (DPT). Male mice were evaluated at 2, 3, 5, 7, and 14 DPT, whereas female mice were evaluated at 2, 7, 10, 14, 18, 26, and 58 DPT.

**Figure 5.**
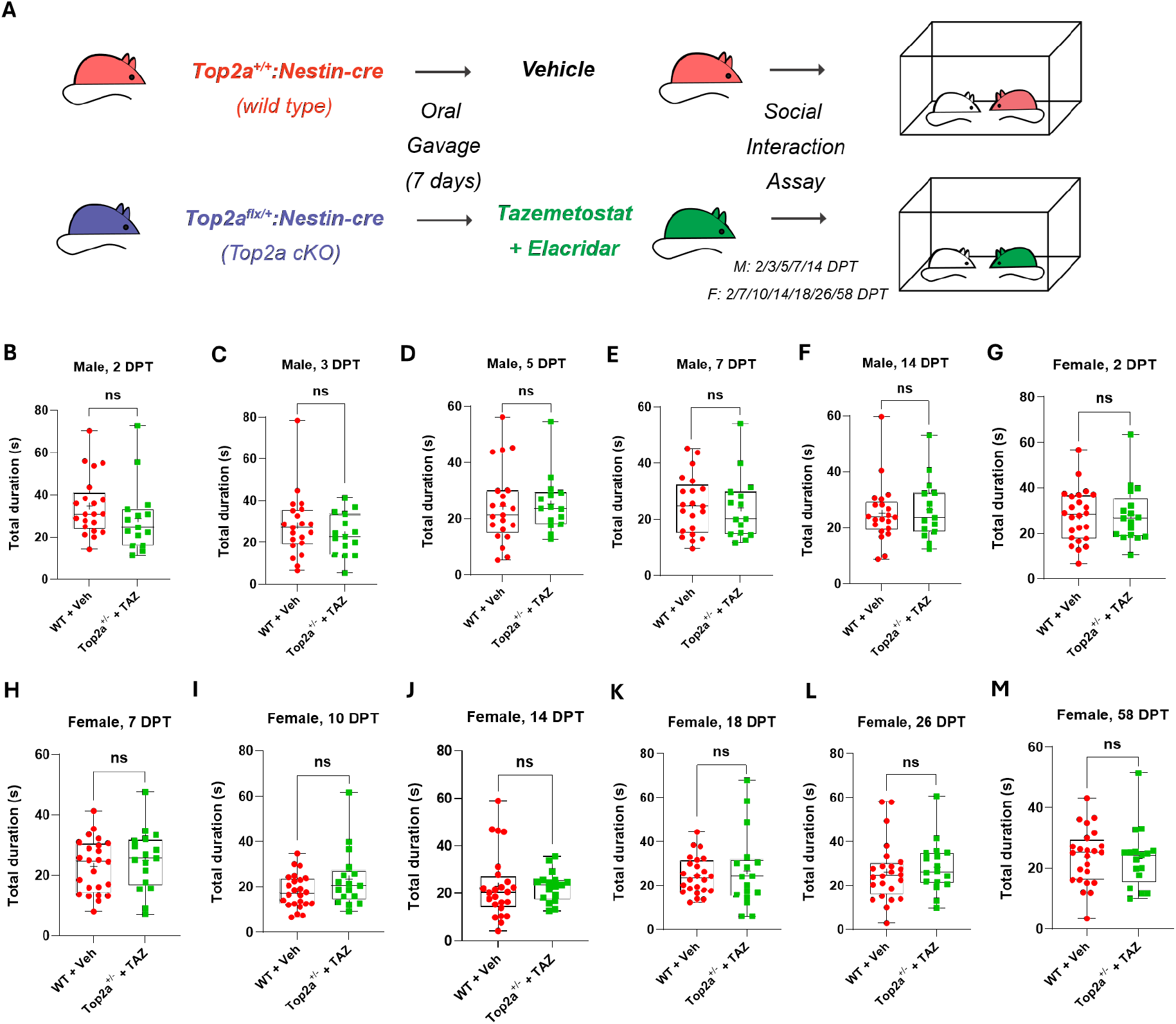
Pharmacological inhibition of PRC2 rescues social deficits in neuronal Top2a cKO mice with sustained effects. (**A**) Experimental schematic illustrating the pharmacological rescue paradigm. Adult Top2a conditional knockout (cKO) mice and wild-type controls were treated with the EZH2 inhibitor tazemetostat in combination with the dual ABCB1/ABCG2 inhibitor elacridar to enhance brain penetration, followed by longitudinal assessment of social behavior after treatment cessation. (**B-F**) Longitudinal assessment of social interaction in male mice following tazemetostat/elacridar and vehicle treatments. Total social interaction duration remained modestly reduced in Top2a cKO mice rescued by tazemetostat (Top2a^+/-^ + TAZ) compared to wild-type vehicle controls (WT + Veh) at early time points (2 and 3 days post-treatment, DPT; B, C), although the reduction was not statistically significant. Rescue mice progressively recovered beginning at 5 DPT (D) and continued to demonstrate social behavior comparable to wild-type levels at 7 and 14 DPT (E, F). (**G-M**) Longitudinal assessment of social interaction in female mice following tazemetostat/elacridar and vehicle treatments. Tazemetostat-treated female Top2a cKO mice (Top2a^+/-^ + TAZ) exhibited robust and sustained normalization of total social interaction duration across all assessed time points compared to wild-type vehicle controls (WT + Veh), persisting throughout the evaluation period from 2 to 58 DPT. Social interaction levels in treated cKO mice were indistinguishable from, and in some cases slightly exceeded, those of wild-type controls throughout the extended examination period. Each dot represents an individual mouse. Box plots indicate median (center line), interquartile range (box), and range (whiskers). All behavioral scoring was performed blinded to genotype. Sample sizes were as follows: male wild-type, *n* = 21; male Top2a cKO, *n* = 15; female wild-type, *n* = 24; female Top2a cKO, *n* = 17. Statistical significance was determined using two-tailed Student’s *t* tests. ns: not significant.

In male mice, tazemetostat treatment robustly restored social interaction in Top2a cKO animals to levels comparable to wild-type controls (Fig. 5B-5F & Supplementary Fig. 1). At 2 and 3 DPT, the mean values for total interaction duration (Fig. 5B & 5C), mean interaction duration (Supplementary Fig. 1A & 1B), and total number of interactions (Supplementary Fig. 1F & 1G) in Top2a cKO mice remained lower than those of wild-type controls. However, beginning at 5 DPT, these measures progressively converged with, and in some cases exceeded, wild-type levels (Fig. 5D-5F; Supplementary Fig. 1C-1E & 1H-1J). Strikingly, this rescue effect persisted for at least 14 days after the final dose, well beyond the expected pharmacokinetic window of the drugs^22-24^.

Encouraged by the persistent rescue effect observed in male mice, we adopted a longer evaluation period for female Top2a cKO mice and observed an even more pronounced durability of rescue. Following the same one-week dosing regimen, tazemetostat-treated female cKO mice exhibited sustained normalization of social interaction that persisted for at least 58 days after treatment cessation (Fig. 5G-5M & Supplementary Fig. 2). Throughout this extended follow-up period, social behavior in treated female cKO mice remained indistinguishable from, and in some cases slightly exceeded, that of wild-type controls for all measured parameters of social interaction, including total interaction duration (Fig. 5G-5M), mean interaction duration (Supplementary Fig. 2A-2G), and total number of interactions (Supplementary Fig. 2H-2N).

To facilitate visualization of the rescue trend, we calculated a rescue index defined as the ratio of mean values between Top2a cKO and wild-type mice at a given time point after treatment (Top2a cKO mean at *X* DPT / WT mean at *X* DPT), normalized to the corresponding ratio measured at baseline prior to treatment (baseline Top2a cKO mean / baseline WT mean). A rescue index value exceeding 1 indicates improved social behavior relative to baseline deficits. In male mice, the rescue index for total social interaction duration increased progressively following treatment cessation, reaching near-complete normalization by 5 days post-treatment (DPT) and remaining stable through 14 DPT (Fig. 6A). Similar trends were observed for mean interaction duration and total number of interactions, indicating coordinated restoration of multiple dimensions of social behavior (Fig. 6B & 6C). These results demonstrate that a brief, 7-day oral PRC2 inhibition regimen is sufficient to induce a sustained rescue of social deficits in male Top2a cKO mice lasting at least two weeks beyond drug withdrawal.

**Figure 6.**
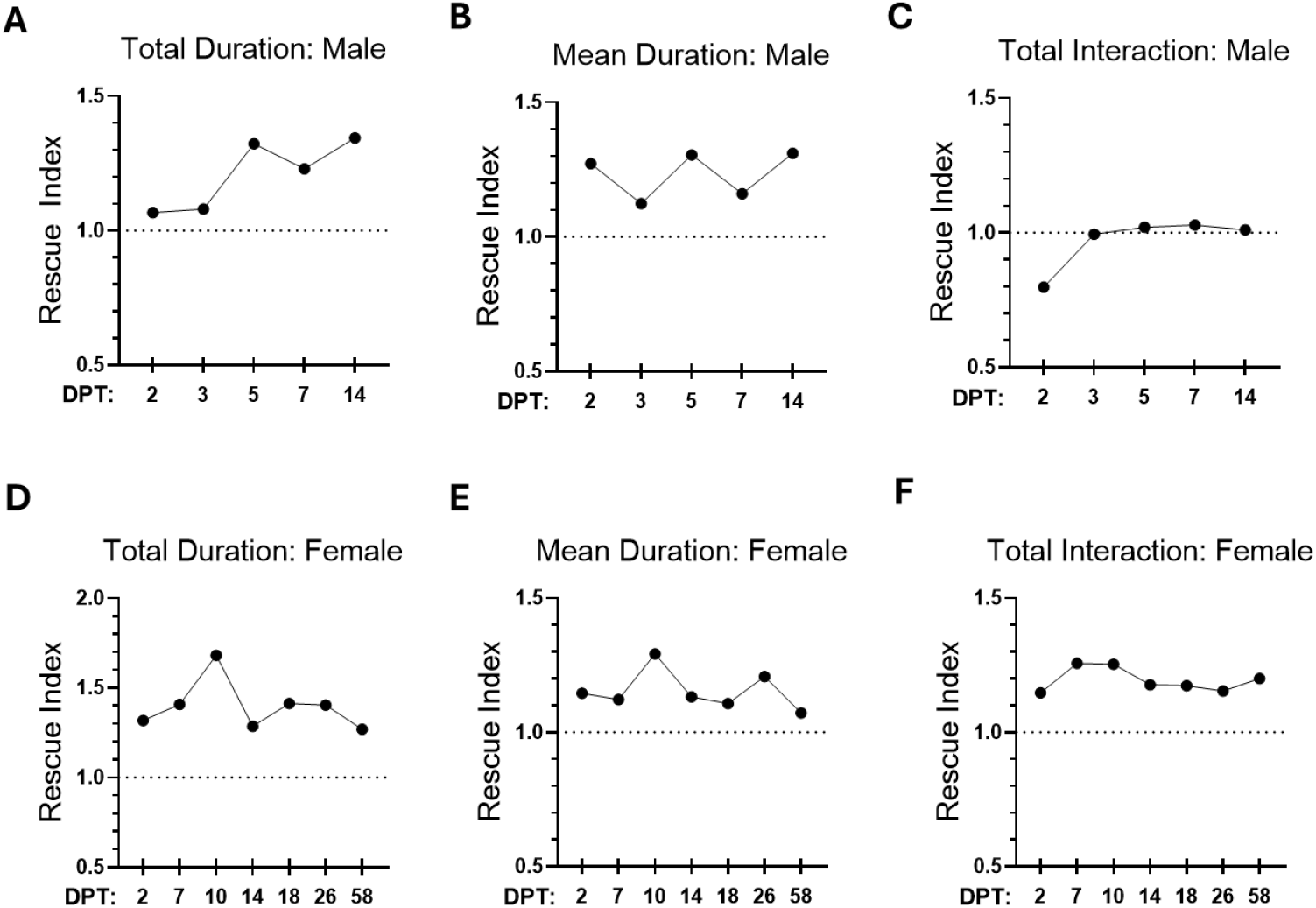
Rescue index analysis reveals durable, sex-dependent restoration of social behavior following transient PRC2 inhibition. (**A-C**) Rescue indices in male mice. Rescue indices were calculated to visualize the magnitude and durability of social behavior rescue following pharmacological PRC2 inhibition. The rescue index is defined as the ratio of mean values between Top2a conditional knockout (cKO) and wild-type mice at a given time point after treatment (Top2a cKO mean at *X* days post-treatment [DPT] / WT mean at *X* DPT), normalized to the corresponding ratio measured at baseline prior to treatment. A rescue index value greater than 1 indicates improvement relative to baseline social deficits. The rescue index for total social interaction duration (A) increased progressively following treatment cessation, reaching near-complete normalization by 5 DPT and remaining stable through 14 DPT. Similar trends were observed for mean interaction duration (B) and total number of interactions (C), indicating coordinated restoration across multiple dimensions of social behavior. (**D-F**) Rescue indices in female mice. The rescue index for total social interaction duration (D) increased rapidly by 2 DPT and remained stably elevated across all subsequent time points, with a transient peak at 10 DPT and persistence through the final assessment at 58 DPT. Comparable temporal dynamics were observed for mean interaction duration (E), whereas the rescue index for total number of interactions (F) remained consistently elevated without any obvious peak.

Strikingly, female Top2a cKO mice exhibited an even more prolonged and robust rescue effect. The rescue index for total social interaction duration rapidly increased to approximately 1.3 by 2 DPT and remained stably elevated across all subsequent time points, with a transient peak reaching 1.68 at 10 DPT, persisting through the final assessment at 58 DPT (Fig. 6D). A similar temporal dynamic was observed for the rescue index of mean interaction duration (Fig. 6E), whereas the rescue index for total number of interactions remained consistently elevated throughout the measurement period without exhibiting a pronounced peak (Fig. 6F).

Importantly, this sustained rescue was observed long after clearance of both tazemetostat^22,23^ and elacridar^24^, suggesting that the behavioral normalization reflects stable epigenetic reprogramming rather than acute neuromodulatory effects. Together, these findings demonstrate that transient inhibition of PRC2 is sufficient to durably reverse genetically induced social deficits caused by neuronal Top2a haploinsufficiency, with effects persisting for weeks to months in a sex-dependent manner.

## DISCUSSION

In this study, we provide direct genetic and pharmacological evidence that neuronal Top2a is a selective and essential regulator of social behavior in mammals and that epigenetic modulation of the Top2a-PRC2 pathway can durably reverse genetically encoded social deficits. By combining a neuron-specific conditional knockout strategy with longitudinal pharmacological rescue experiments, our work establishes the neuronal Top2a-PRC2 axis as a conserved, targetable epigenetic pathway governing social behavior, while also revealing unexpected specificity of behavioral regulation and remarkable durability of the rescue effect.

Our findings provide critical genetic validation of a Top2a-PRC2 pathway previously identified largely through pharmacological approaches^11^. Prior work demonstrated that Top2a inhibition disrupts social behavior through aberrant PRC2-mediated H3K27 trimethylation and that PRC2 inhibition rescues social deficits in zebrafish. Here, we extend these observations to mammals by showing that Top2a haploinsufficiency in neurons successfully recreated the social deficit phenotype observed previously through chemical inhibition of Top2a^11^. These findings verify the Top2a-PRC2 pathway as a genuine molecular mechanism required for the normal development of mammalian social behavior rather than a pharmacological artifact.

Another key finding of this study is that neuron-specific Top2a haploinsufficiency selectively impairs social interaction without inducing restricted and repetitive behaviors, cognitive deficits, anxiety-like behavior, or depressive-like phenotypes. This dissociation is surprising given that ASD is typically characterized by concurrent impairments in social behavior and restricted and repetitive behaviors, often accompanied by additional behavioral comorbidities mentioned above, a phenomenon frequently recapitulated by mouse models of ASD risk gene knockout^25-32^. Our results indicate that disruption of social behavior can occur independently of other ASD-relevant phenotypes and that neuronal Top2a plays a specialized role in regulating sociality.

This phenomenon contrasts with our prior chemical inhibition study^11^, in which embryonic exposure to Top2 inhibitors induced not only social deficits but also restricted and repetitive behaviors in mice. One likely explanation for this discrepancy is cell-type specificity. While the Nestin-Cre strategy used in this study specifically targets neurons, systemic pharmacological inhibition of Top2a during development as implemented in our previous work^11^ likely affects multiple cell populations within the brain. This led us to hypothesize that restricted and repetitive behaviors induced by Top2a inhibition may be mediated by non-neuronal cell types.

Both astrocytes and microglia represent compelling candidates in this context. A growing body of literature has implicated these cell types in the regulation of repetitive and compulsive behaviors^33-38^. For example, a molecularly defined population of Crym-positive astrocytes in the striatum has been shown to gate perseverative behavior by regulating corticostriatal synaptic transmission, with astrocyte-specific loss of Crym inducing robust perseveration through altered excitatory-inhibitory balance in striatal circuits^34^.

In parallel, a series of seminal studies has established a central role for microglia in the control of pathological grooming and other compulsive behaviors through the Hoxb8 pathway. Initial genetic analyses revealed that *Hoxb8* disruption leads to highly penetrant compulsive grooming and hair removal in mice^33^. Subsequent lineage-tracing and bone marrow transplantation experiments demonstrated that, within the brain, *Hoxb8* expression is restricted to a bone marrow-derived microglial population, and that replacement of mutant microglia with wild-type counterparts is sufficient to rescue pathological grooming behavior^36^. Defective Hoxb8 microglia induced profound corticostriatal circuit abnormalities, linking microglial genetic perturbation to circuit-level dysfunction classically associated with obsessive-compulsive disorder and autism spectrum disorders. Notably, Hoxb8 mutant mice also exhibit social behavioral impairments and anxiety-like phenotypes^38^. Finally, optogenetic manipulation of Hoxb8 microglia provided causal, region-specific evidence that microglial activity is sufficient to acutely drive grooming and anxiety behaviors. Stimulation of Hoxb8 microglia within the dorsomedial striatum or medial prefrontal cortex selectively induced grooming, whereas stimulation in amygdala nuclei elicited anxiety-like behavior^37^.

These prior findings suggest that Top2a-dependent mechanisms in non-neuronal cell types, particularly astrocytes and microglia, may preferentially regulate restricted and repetitive behavioral domains, whereas neuronal Top2a is selectively required for social behavior regulation. Given the essential roles of Top2a in transcriptional regulation^39^ and chromatin organization^10,40^, epigenetic dysregulation in glial populations could alter circuit maturation or synaptic homeostasis in ways that specifically bias toward repetitive or compulsive phenotypes. Future studies employing microglia- or astrocyte-specific Top2a deletion will be critical for testing this hypothesis and for dissecting how cell-type-specific epigenetic programs independently govern distinct ASD-relevant behavioral domains.

Perhaps the most striking finding of this study is the durability of behavioral rescue following transient PRC2 inhibition. A single, one-week oral dosing regimen of tazemetostat produced social behavior rescue lasting at least two weeks in males and two months in females, far exceeding the temporal window typically observed for neuromodulatory drugs targeting neurotransmitter systems^17,18^. Such persistence strongly suggests that PRC2 inhibition induces stable changes in gene regulatory states rather than transient modulation of synaptic signaling. This durability also supports a model in which epigenetic interventions can reset maladaptive transcriptional programs underlying social behavior deficits. In contrast to conventional neuropsychiatric drugs that require continuous administration to maintain efficacy, our results suggest that epigenetic modulators may offer the possibility of long-lasting therapeutic benefits following a short-term treatment regimen. The sex-dependent differences in rescue magnitude and durability observed here raise intriguing questions regarding sex-specific epigenetic stability, chromatin dynamics, and circuit plasticity that warrant further investigation. Notably, male and female mice were assessed over different longitudinal time windows in the present study; therefore, a direct, side-by-side comparison using matched testing durations will be necessary to rigorously evaluate sex differences in the persistence of rescue effects.

Our findings have broad implications for understanding and treating neurodevelopmental and psychiatric disorders characterized by social dysfunction. First, they provide strong evidence that social behavior is governed by distinct molecular and epigenetic mechanisms that can be genetically and pharmacologically dissociated from other behavioral domains. Second, they establish epigenetic modulation—specifically targeting the Top2a-PRC2 axis—as a viable strategy for inducing durable behavioral recovery in a mammalian system. From a translational perspective, the use of tazemetostat, a clinically approved EZH2 inhibitor, demonstrates the therapeutic relevance of our findings. While additional work is needed to assess safety, optimal dosing, and developmental timing, our results suggest that targeting epigenetic regulators may offer fundamentally different and potentially more durable therapeutic outcomes than existing neuromodulatory approaches.

In summary, this study demonstrates that neuronal Top2a is genetically required for normal social behavior in mice and that transient inhibition of PRC2 can durably rescue social deficits caused by Top2a deficiency. These findings establish the neuronal Top2a-PRC2 axis as a conserved epigenetic pathway regulating social behavior and showcase epigenetic modulation as a promising avenue for treating core symptoms of ASD and related neuropsychiatric disorders. Future studies dissecting cell-type-specific roles of Top2a and PRC2, particularly in glial populations, will be essential for fully understanding how distinct ASD-relevant behaviors are differentially regulated and for refining epigenetic therapeutic strategies.

## MATERIALS AND METHODS

### Mouse husbandry

The Nes-cre mouse line on a C57BL/6J background was obtained from The Jackson Laboratory (strain no. 003771). The Top2a^tm1a(KOMP)Wtsi^ conditional knockout mouse line was rederived from embryonic stem cells generated by the Knockout Mouse Project (KOMP) and obtained through the Mutant Mouse Resource & Research Centers (MMRRC; stock no. 064016-UCD). Rederivation was performed by the University of Utah Transgenic and Gene Targeting Core onto a C57BL/6N background. To generate experimental animals, male and female Top2a^flx/+^ mice were crossed with homozygous Nestin-Cre mice. Offspring were genotyped by PCR prior to weaning to determine the presence of the floxed Top2a allele and the Nestin-Cre transgene. Two experimental groups were established: Top2a^flx/+^;Nestin-Cre mice (conditional knockout, cKO) and Top2a^+/+^;Nestin-Cre littermates (wild-type controls). Genotypes were reconfirmed by PCR prior to enrollment in behavioral and pharmacological rescue experiments, and only mice with verified genotypes were included in analyses. C57BL/6NJ mice used as social stimulus animals were obtained from The Jackson Laboratory (strain no. 005304) and bred in parallel with experimental cohorts to generate age-matched stimulus animals for social interaction assays.

Mice were housed in groups of up to five per cage under controlled environmental conditions (20-26 °C; ∼50% relative humidity) on a 12 h light/12 h dark cycle. Standard chow and water were provided *ad libitum*, except during behavioral testing. Animals were allowed to acclimate to the procedure room prior to testing, and all behavioral assays were conducted during the light phase of the circadian cycle. All animal procedures were approved by the University of Washington Institutional Animal Care and Use Committee (IACUC) and were performed in accordance with applicable federal animal welfare regulations.

### Tail tissue lysate and Genotyping

Genomic DNA was prepared from mouse tail tissue using the HotSHOT alkaline lysis method. Briefly, 1-2 mm tail biopsies were incubated in alkaline lysis buffer (25 mM NaOH, 0.2 mM EDTA) at 95 °C for 1 h to release genomic DNA, followed by neutralization with 40 mM Tris-HCl. The resulting lysate was mixed thoroughly and used directly as PCR template without further purification. Top2a wild-type and floxed alleles, as well as the Nestin-Cre transgene, were identified by PCR using established primer sets. Primer sequences, PCR conditions, and expected amplicon sizes are provided in Supplementary Table 1.

### Social interaction assay

Social interaction assays were conducted as previously described with minor modifications^11^. Mice were initially tested at 6 months of age and were retested at 14-16 months of age following the rescue experiment. Each test mouse was introduced into an unfamiliar, neutral Allentown cage and allowed to acclimate for 2 min. An unfamiliar, weight-, sex-, and age-matched wild-type conspecific from a different litter was then introduced as the social stimulus. To distinguish the test mouse from the stimulus animal, the test mouse was marked with an odorless, non-toxic colored marker. Social interaction sessions lasted 5 min and were recorded in their entirety using a Raspberry Pi Camera Module connected to a Raspberry Pi. Social behaviors were scored offline using BORIS^41^ by experimenters blinded to genotype. Social interaction was defined as periods during which the test mouse actively investigated the stimulus mouse, including sniffing of the facial, abdominal, or anogenital regions, grooming, or closely following the stimulus mouse during exploration of the cage. Investigation initiated by the stimulus mouse toward the test mouse was not scored. All female mice were tested during the metestrus and diestrus stages of the estrous cycle to minimize behavioral variability.

### Novel object exploration assay

Novel object exploration assays were conducted as previously described^11^ using mice aged 6-7 months. Female mice were tested during the metestrus and diestrus stages of the estrous cycle. The assay was performed in an unfamiliar, neutral Allentown cage using the same experimental setup as the social interaction assay. Following an acclimation period, a wooden block was introduced into the cage as the novel object. The object was fully wrapped with tape during each trial, and the tape was removed and replaced between tests to eliminate residual odor cues. The total duration of object investigation over a 5-min period was quantified using BORIS.

### Open field assay

The open field assay was conducted in a rectangular arena consisting of an outer white cardboard enclosure and an inner transparent plexiglass chamber (60 × 48 cm), supported by a black frame and enclosed by four white walls (40 cm high). The arena floor was divided into two zones of equal area: a central rectangular zone (42.43 × 33.94 cm) and a surrounding peripheral zone defined as the area within 8.79 cm and 7.03 cm from the walls. At the start of each trial, mice were placed in the center zone, and behavior was recorded for 10 min using a Raspberry Pi Camera Module. Locomotor activity was tracked and analyzed using AnyMaze behavioral tracking software (version 7.33). The percentage of time spent in the center zone and average velocity were quantified as measures of anxiety-like behavior and general locomotor activity, respectively. Male mice were tested at 6 months of age, and female mice were tested at 8 months of age during the metestrus and diestrus stages of the estrous cycle.

### Novel object recognition assay

Novel object recognition assays were conducted following a published protocol^42^ using mice aged 7-8 months. Female mice were tested during the metestrus and diestrus stages of the estrous cycle. The assay was performed in the same arena used for the open field assay. Each mouse was first habituated to the arena for 5 min. Following habituation, the mouse was removed, and two identical cubicle wooden blocks were placed in opposite corners of the arena, each positioned 7 cm from the side walls. The mouse was then reintroduced into the center of the arena and allowed to explore the arena and the two objects for 10 min. Six hours later, one of the familiar objects was replaced with a novel object of similar size but distinct shape (cylindrical) and color. The mouse was returned to the arena and allowed to explore both objects for an additional 10 min. Mouse behavior was recorded, and the total duration of object investigation was quantified using BORIS. Recognition memory was assessed by calculating the recognition index, defined as the time spent investigating the novel object divided by the sum of time spent investigating the novel and familiar objects.

### Marble burying assay

The marble burying assay was performed in an unfamiliar, neutral Allentown cage filled with 4.5 cm of fresh standard corncob bedding. Twenty black glass marbles were gently placed on the bedding surface in a uniform 4 × 5 arrangement with equal spacing. Each test mouse was introduced into the bottom-right corner of the cage and allowed to explore freely for 30 min. At the end of the trial, both the number of displaced marbles and the number of marbles covered by more than 50% of their surface area with bedding were scored. Photographic records were obtained for documentation. Male mice were tested at 7 months of age, and female mice were tested at 7-9 months of age during the metestrus and diestrus stages of the estrous cycle.

### Tail suspension assay

The tail suspension assay was conducted following a published protocol^42^. Male mice were tested at 7 months of age, and female mice were tested at 9 months of age during the metestrus and diestrus stages of the estrous cycle. Briefly, each mouse was suspended from the rail of a tail suspension frame using adhesive tape placed 2 cm from the tip of the tail, positioning the animal approximately 60 cm above the surface of a table. Following a 2-min acclimation period, behavior was recorded for 5 min using a Raspberry Pi Camera Module. Immobility duration was scored using BORIS and was defined as periods during which the mouse hung passively and remained completely motionless.

### T-maze assay

The T-maze assay was conducted using a T-shaped tubular apparatus equipped with movable doors separating the start compartment from the two choice arms. Each mouse completed eight consecutive trials per session. At the beginning of each trial, the mouse was placed in the start arm and confined for 1 min before the first trial or 15 s for subsequent trials, after which the start door was opened to allow free exploration of the maze. A choice was recorded when the mouse fully entered one of the arms, defined as the placement of all four paws within the selected arm. The corresponding door was then closed to confine the mouse in that arm for 30 s. Between trials, mice were removed from the apparatus, and the maze was cleaned with 70% ethanol and dried to eliminate residual odor cues. Mice were tested at 9 months of age; female mice were tested during the metestrus and diestrus stages of the estrous cycle.

### Elacridar and Tazemetostat formulation

Elacridar, tazemetostat, and their corresponding vehicle solutions were prepared fresh daily prior to oral administration. Elacridar and tazemetostat were purchased from BOC Sciences (catalog nos. B0084-311800 and B0084-462339, respectively). Elacridar was prepared as a 5 mg/mL solution in a DMSO:Cremophor EL:water vehicle (1:2:7, v/v/v). Elacridar powder was first dissolved in DMSO at 50 mg/mL by vortexing and heating in boiling water for 10 min. Two parts Cremophor EL (MedChemExpress; cat. no. HY-Y1890) and seven parts water were then added sequentially to achieve the final formulation. Tazemetostat was prepared as a 10 mg/mL suspension in water containing 0.5% (v/v) hydroxypropyl methylcellulose (HPMC; Thermo Scientific; CAS 9004-65-3) and 0.1% (v/v) Tween-80 (MedChemExpress; cat. no. HY-Y1891). The mixture was vortexed and sonicated for 10 min to ensure uniform suspension and resuspended immediately before each oral administration.

### Oral administration of Elacridar and Tazemetostat

Top2a conditional knockout mice and wild-type littermates aged 14-15 months were subjected to a 7-day oral dosing regimen via oral gavage. All mice were weighed prior to drug administration, and dosing volumes were calculated based on body weight. Elacridar and tazemetostat were administered sequentially each day. On the first day of treatment, mice received elacridar (100 mg/kg, oral gavage), followed by tazemetostat (100 mg/kg, oral gavage) after a 3-h interval. On treatment days 2-7, mice received elacridar (50 mg/kg, oral gavage), followed by tazemetostat (100 mg/kg, oral gavage) after a 2-h interval. Wild-type control mice received the corresponding vehicle solutions following the same administration schedule and timing. Social interaction assays were performed after completion of the 7-day dosing regimen.

### Statistical analysis

All graphs were generated using GraphPad Prism. Statistical analyses were performed using Student’s *t* tests. Two-tailed *t* tests were used unless otherwise specified. For a priori hypotheses testing reduced social behavior in conditional knockout mice relative to wild-type controls, one-tailed *t* tests were applied. A *P* value < 0.05 was considered statistically significant.

## ACKNOWLEDGEMENTS

We thank Dr. Olaf van Tellingen at The Netherlands Cancer Institute for valuable discussion and suggestions on elacridar/tazemetostat formulation and delivery. We thank the University of Washington Office of Comparative Medicine for providing mouse husbandry support. This work was supported by the National Institute of Environmental Health Sciences (NIEHS) of the NIH under the award number R00ES031050. The content is solely the responsibility of the authors and does not necessarily represent the official views of the NIH.

## AUTHOR CONTRIBUTIONS

B.H., Y.M., and C.H. conducted experiments and analyzed data. Y.G. conceived and designed the study, interpreted the data, and wrote the manuscript.

## DATA AND MATERIALS AVAILABILITY

All data needed to evaluate the conclusions in the paper are present in the paper and/or the Supplementary Materials.

