## Supplementary Materials for "Top2a-dependent neuronal regulation of social behavior and persistent rescue of social deficit through PRC2-mediated epigenetic reprogramming"

**This PDF file includes:**

Supplementary Figures 1-2

Supplementary Table 1

### SUPPLEMENTARY FIGURE

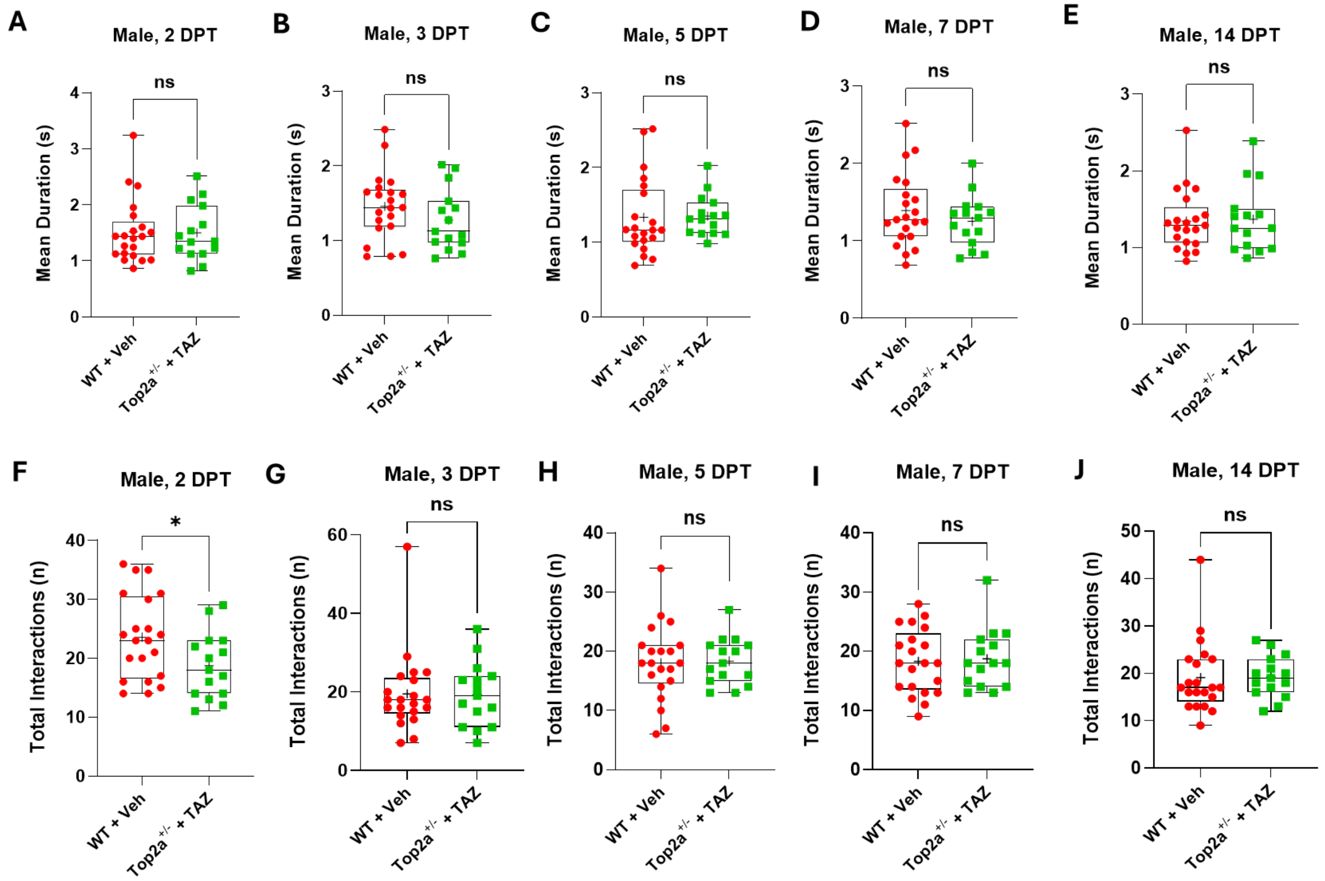


**Supplementary Figure 1. Longitudinal analysis of mean interaction duration and total interaction number in male mice following PRC2 inhibition.**

(**A-E**) Mean duration of social interactions in male mice at 2, 3, 5, 7, and 14 days post-treatment (DPT), respectively. At early time points (2 and 3 DPT), tazemetostat-rescued Top2a conditional knockout (cKO) mice (Top2a$^{+/-}$ + TAZ) continued to exhibit slightly reduced (not statistically significant) mean interaction duration compared to wild-type vehicle controls (WT + Veh) (A, B). Beginning at 5 DPT, mean interaction duration in treated Top2a cKO mice progressively converged with, and in some cases exceeded, wild-type levels (C-E).

(**F-J**) Total number of social interactions in male mice at 2, 3, 5, 7, and 14 DPT, respectively. Total interaction number remained reduced at 2 DPT (F) but recovered beginning at 3 DPT and remained comparable to wild-type levels through 14 DPT (G-J).

Each dot represents an individual mouse. Box plots indicate median (center line), interquartile range (box), and range (whiskers). All behavioral scoring was performed blinded to genotype. Sample sizes were as follows: male wild-type, $n$ = 21; male Top2a cKO, $n$ = 15; female wild-type, $n$ = 24; female Top2a cKO, $n$ = 17. Statistical significance was determined using two-tailed Student’s $t$ tests. ns: not significant.

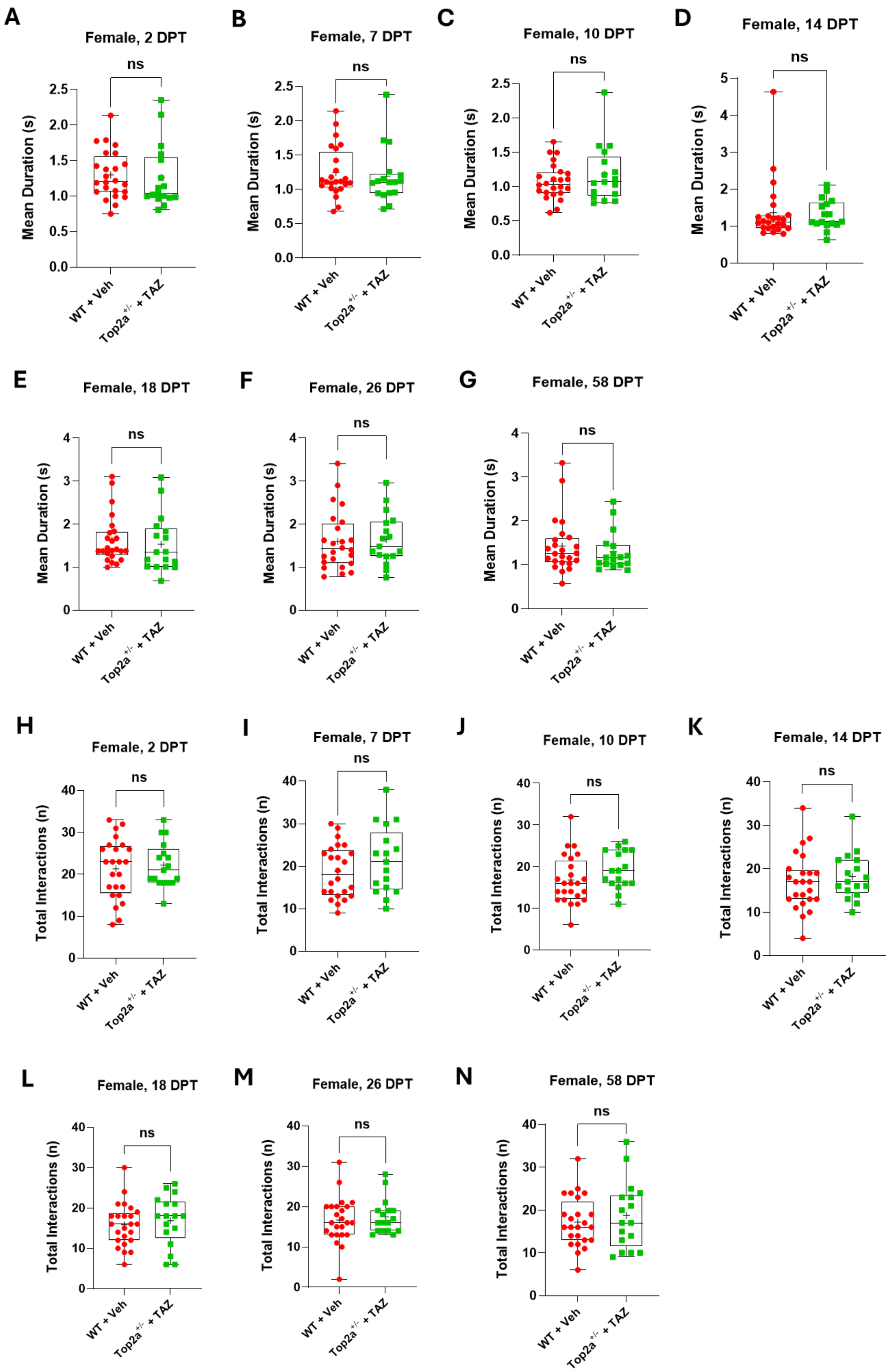
A
Female, 2 DPT
B
Female, 7 DPT
C
Female, 10 DPT
D
Female, 14 DPT
E
Female, 18 DPT
F
Female, 26 DPT
G
Female, 58 DPT
H
Female, 2 DPT
I
Female, 7 DPT
J
Female, 10 DPT
K
Female, 14 DPT
L
Female, 18 DPT
M
Female, 26 DPT
N
Female, 58 DPT
ns
Mean Duration (s)
Total Interactions (n)
WT + Veh
Top2a+/- + TAZ

**Supplementary Figure 2. Sustained rescue of social behavior parameters in female mice following PRC2 inhibition.**

(**A-G**) Mean duration of social interactions in female mice at 2, 7, 10, 14, 18, 26, and 58 days post-treatment (DPT), respectively. Tazemetostat-treated Top2a conditional knockout (cKO) mice (Top2a$^{+/-}$ + TAZ) exhibited sustained normalization of mean interaction duration across all assessed time points, remaining indistinguishable from or exceeding wild-type vehicle control (WT + Veh) levels throughout the extended examination period.

(**H-N**) Total number of social interactions in female mice at the same time points. Total interaction number in tazemetostat-treated Top2a cKO mice (Top2a$^{+/-}$ + TAZ) remained consistently elevated relative to baseline deficits and comparable to wild-type vehicle controls (WT + Veh) across the entire 58-day observation window.

Each dot represents an individual mouse. Box plots indicate median (center line), interquartile range (box), and range (whiskers). All behavioral scoring was performed blinded to genotype. Sample sizes were as follows: male wild-type, $n$ = 21; male Top2a cKO, $n$ = 15; female wild-type, $n$ = 24; female Top2a cKO, $n$ = 17. Statistical significance was determined using two-tailed Student's $t$ tests. ns: not significant.

### SUPPLEMENTARY TABLE

**Supplementary Table 1. Primer sequences, PCR conditions, and expected amplicon sizes for genotyping.**

The following protocol and primer design are established by Mutant Mouse Resource & Research Center at UC Davis (MMRRC: 064016-UCD) and the Jackson Laboratory (Protocol: 31758).

PCR protocol:

| Reagent/Constituent | Volume (µL) |
|---|---|
| 2X Taq Polymerase (NEB M0270) | 10 |
| Forward Primer | 2 |
| Reverse Primer | 2 |
| Water | 4 |
| DNA | 2 |
| **TOTAL VOLUME OF REACTION** | 20 |

PCR condition:

| Steps | Temp (ºC) | Time (m:s) | # of Cycles |
|---|---|---|---|
| 1. Initiation/Melting | 95 | (10:00) | 1 |
| 2. Denaturation | 95 | (00:30) | 40X |
| 3. Annealing | 54 | (00:30) | |
| 4. Elongation | 72 | (00:45) | |
| 5. Amplification | 72 | (10:00) | 1 |
| 6. Finish | 4 | ∞ | 1 |

Primer sequences:

| Primers | Nucleotide Sequence (5' - 3') |
|---|---|
| 1 | GAGATGGCGCAACGCAATTAATG |
| 2 | CAAGCTGGGTAAGATTGGTGAACC |
| 3 | CTTACAGGTTACACAGGCAGTTAGC |
| 4 | CATTTGAGAACGGTAACCTGGATGG |
| 5 | TTGCTAAAGCGCTACATAGGA |
| 6 | GCCTTATTGTGGAAGGACTG |
| 7 | CCTTCCTGAAGCAGTAGAGCA |

Primer combinations:

| Primer Combination | Band (bp) | Genotype |
|---|---|---|
| 1 & 2 | 319 | Top2a floxed |
| 3 & 4 | 285 | Top2a Wildtype |
| 6 & 7 | 150 | Nes-Cre Mutant |
| 5 & 6 | 246 | Nes-Cre Wildtype |
